# Bidirectional intraflagellar transport is restricted to two sets of microtubule doublets in the trypanosome flagellum

**DOI:** 10.1101/329300

**Authors:** Eloïse Bertiaux, Adeline Mallet, Cécile Fort, Thierry Blisnick, Serge Bonnefoy, Jamin Jung, Moara Lemos, Sergio Marco, Sue Vaughan, Sylvain Trépout, Jean-Yves Tinevez, Philippe Bastin

## Abstract

Intraflagellar transport (IFT) is the rapid bidirectional movement of large protein complexes driven by kinesin and dynein motors along microtubule doublets of cilia and flagella. Here we used a combination of high-resolution electron and light microscopy to investigate how and where these IFT trains move within the flagellum of the protist *Trypanosoma brucei*. Focused Ion Beam Scanning Electron Microscopy (FIB-SEM) analysis of trypanosomes showed that trains are found almost exclusively along **two sets of doublets (3-4 and 7-8)** and distribute in two categories according to their length. High-resolution live imaging of cells expressing mNeonGreen::IFT81 or GFP::IFT52 revealed for the first time IFT trafficking on two **parallel lines within the flagellum. Anterograde and retrograde IFT occur on each of these lines**. At the distal end, a large individual anterograde IFT train is converted in several smaller retrograde trains in the space of 3-4 seconds while remaining on the same **side of the axoneme**.

## Introduction

Intraflagellar transport (IFT) is the movement of molecular motors and multi-protein complexes that carry tubulin and other flagellar components to the tip of cilia and flagella for assembly (Craft et al., 2015; Kozminski et al., 1993). One or more kinesin motors are responsible for anterograde transport whereas a dynein motor returns the trains to the base during retrograde transport (Prevo et al., 2017). These moving protein complexes have been termed IFT trains (Pigino et al., 2009). Absence of IFT prevents construction of most cilia and flagella whereas perturbation of IFT components can impact on the structure and function of the organelle, as observed in multiple human genetic diseases (Beales et al., 2007; Dagoneau et al., 2009; Halbritter et al., 2013; Perrault et al., 2012).

Quantification of IFT in animal cells, green algae, trypanosomes or ciliates revealed remarkably high speed (0.5-5 µm per second) and frequency (∼1-3 trains per s) of IFT trains in both directions (Besschetnova et al., 2010; Brooks and Wallingford, 2012; Buisson et al., 2013; Iomini et al., 2001; Prevo et al., 2015; Snow et al., 2004; Wheeler et al., 2015; Williams et al., 2014; Wingfield et al., 2017). When anterograde trains reach the distal end, they are converted in usually smaller retrograde trains in the space of a few seconds (Buisson et al., 2013; Chien et al., 2017; Mijalkovic et al., 2017). Trains are fairly large complexes of >20 proteins (Taschner and Lorentzen, 2016) associated to molecular motors whose size is above 1 megadalton (Rompolas et al., 2007), raising the question of how they are organised within the flagellum during anterograde and retrograde trafficking.

In transmission electron microscopy (TEM), trains appear as electron dense particles sandwiched between microtubule doublets and the flagellum membrane. This was visualised in only two species so far: the green algae *Chlamydomonas reinhardtii* (Pigino et al., 2009; Ringo, 1967; Vannuccini et al., 2016) and the protist *Trypanosoma brucei* (Absalon et al., 2008). Recently, an elegant study using correlative light electron microscopy in *Chlamydomonas* cells expressing a fluorescent IFT protein showed that anterograde trains are positioned on the B-tubule of each microtubule doublet whereas retrograde trains are found on the A-tubule (Stepanek and Pigino, 2016). In this organism, IFT trains appeared on all 9 microtubule doublets (Stepanek and Pigino, 2016). By contrast, transmission electron microscopy (TEM) suggested that electron dense particles looking like IFT trains are restricted to microtubule doublets 3-4 and 7-8 of the axoneme in *Trypanosoma brucei* (Fig. 1A)(Absalon et al., 2008; Hoog et al., 2016). The restricted presence of IFT trains on two **sets of doublets** raises two hypotheses. First, a tantalising explanation would be that **some specific doublets** serve as tracks for anterograde transport and **others** for retrograde trafficking, hence allowing independent control of each mode of transport (Fig. 1B, Model 1). Second, each **individual doublet** could serve as a double track for IFT (Fig. 1B, Model 2), as shown in *Chlamydomonas*. In that situation, train frequency and speed could be similar or different between doublets. In both models, once anterograde trains reach the distal end of the flagellum they need to be converted to retrograde trains in order to return back to the proximal end. Does conversion happen on the same doublet that the anterograde train travelled along or does each anterograde train enter a common pool of material at the distal end of the flagellum where retrograde trains are then assembled? Here, we investigated IFT train distribution along the length of the trypanosome flagellum using Focused Ion Beam Scanning Electron Microscopy (FIB-SEM) to get a three dimensional view and to measure the length of each IFT train using a 10nm Z-axis increment. We formally demonstrate that trains are indeed mostly found on doublets 3-4 and 7-8 and that they fall in two categories defined by their length. Using high-resolution live cell imaging, we reveal which hypothesis is correct for the distribution of anterograde and retrograde trafficking and we apply kymograph analysis to examine the conversion of anterograde to retrograde trains.

**Fig. 1.**
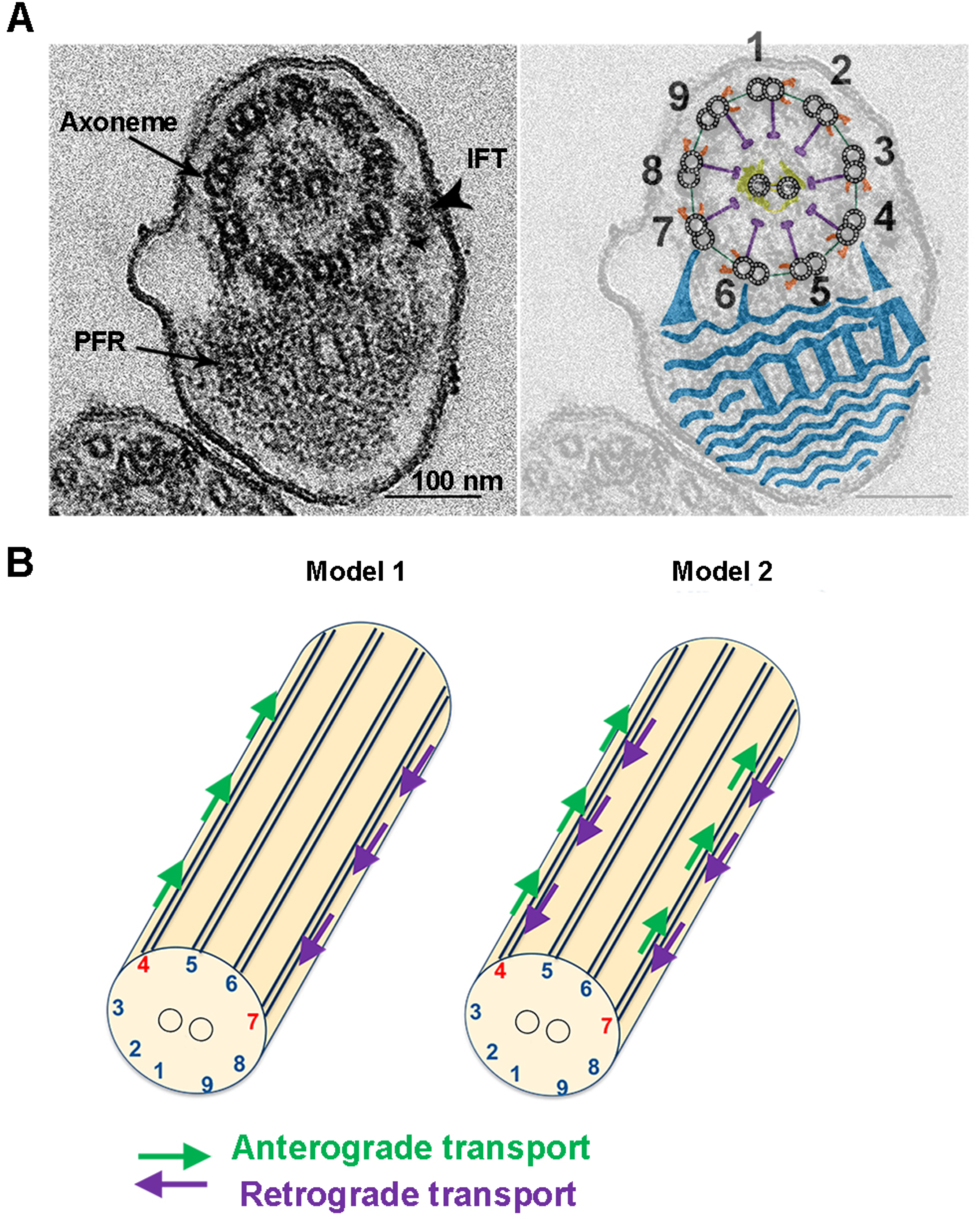
Positioning of IFT trains in the trypanosome flagellum and models for IFT trafficking. **(A)** Cross-section of the trypanosome flagellum observed by conventional TEM. The arrowhead indicates an IFT particle positioned at the level of doublet 4. The cartoon shows the main structural components of the axoneme with the numbering of microtubule doublets (Branche et al., 2006) superposed on the original image. Doublet numbering follows the conventional rules: a line perpendicular to the middle axis of the central pair microtubules is drawn and makes contact with the A-tubule of only one doublet that is defined as number 1. The numbering follows the clockwise orientation defined by the dynein arms. Dynein arms are shown in orange, radial spoke in violet, central pair projections in yellow and the PFR in blue. **(B)** The restricted presence of IFT on doublets 3-4 and 7-8 can be explained by two models: **either some doublets are used specifically for anterograde transport and the other ones support retrograde IFT (Model 1), or bidirectional trafficking takes place on all doublets (Model 2). Only one doublet was highlighted for the sake of clarity.**

## RESULTS

### FIB-SEM analysis revealed the length and 3-D distribution of IFT trains

To obtain a global view of the 3-D distribution of IFT trains by electron microscopy, different approaches are available. Since the twisted shape of the trypanosome flagellum (Sherwin and Gull, 1989) restricts the use of conventional transmission electron tomography to short portions of the axoneme, we turned to FIB-SEM that allows the collection of hundreds of sequential electron microscopy images at 10nm Z-resolution (Kizilyaprak et al., 2014). Wild-type trypanosomes were chemically fixed, dehydrated and embedded in resin in conditions similar to classic transmission electron microscopy. **Acquisition in x-y relies on scanning electron microscopy collecting backscattered electrons and therefore provides a lower resolution compared to transmission electron microscopy. However, invaluable z-information is gained by milling the block face with 10 nm increments using the ion beam. In these conditions, isotropic large volumes can be acquired with ultra-structural resolution at 10nm of voxel size to analyse intracellular structures in long portions of flagella. This turned out to be a considerable advantage to monitor IFT train positioning.** Four samples were reconstructed, each containing several trypanosomes. Navigating through the volume of each sample revealed the typical trypanosome architecture with the cell body containing the subpellicular microtubules, the nucleus, the large mitochondrion and its kinetoplast (mitochondrial DNA), the endoplasmic reticulum, the Golgi apparatus or the glycosomes (Video S1)(Fig. 2A). The shape, size and distribution of these organelles were in agreement with published data based on electron tomography (Lacomble et al., 2009) or serial block face sectioning (Hughes et al., 2017). Flagella were clearly recognised in both cross and longitudinal sections, including multiple cases where the proximal portion was seen to emerge from the flagellar pocket (Video S1). The base of the flagellum displayed the typical organisation with the basal body, the transition zone, the axoneme and finally the axoneme and the paraflagellar rod (PFR), a lattice-like structure (Hughes et al., 2012) essential for motility (Bastin et al., 1998). Flagella were correctly attached to the cell body with the exception of the distal end that is always free as expected (Sherwin and Gull, 1989).

**Fig. 2.**
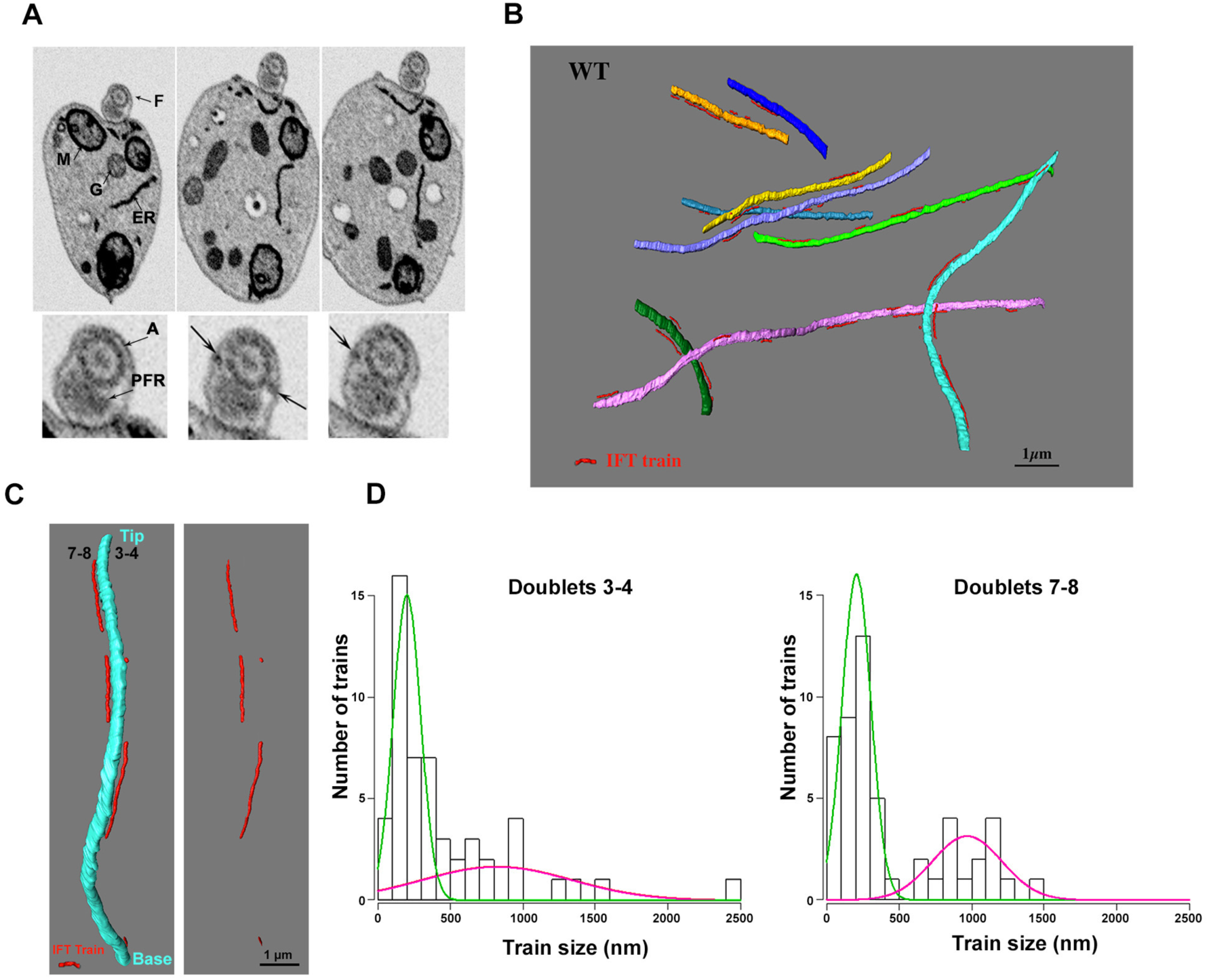
IFT trains of similar length are distributed on doublets 3-4 and 7-8. **(A)** Successive images from Video S1 showing wild-type trypanosomes analysed by FIB-SEM. Each image corresponds to a Z-stack of 3 slices between positions 424 and 475. The progression is from posterior to anterior. The top panels show low magnification of the cell body where major organelles are indicated (ER, endoplasmic reticulum, F, flagellum, G, glycosomes, M, mitochondrion). Bottom: A zoom of the flagellum area is shown with the axoneme (A) and the PFR. The arrows indicate IFT particles. **(B)** Portions of flagella reconstructed after FIB-SEM. Each axoneme is shown with a different colour and IFT trains are in red (for animation, see Video S2). **(C) Another** example of a flagellum from a wild-type trypanosome coming from a different stack than the one presented in A-B with the axoneme (sky blue) and several IFT trains (red). **Doublet numbers and flagellum orientation (BB, basal body and tip) are indicated. (D)** Length of the IFT trains on doublets 4 (green) and 7 (magenta) determined from FIB-SEM analysis. Data are coming from 27 portions of flagella representing a total axoneme length of 166 µm. Two populations can be separated with short trains (green) and longer ones (magenta, see text for details).

Importantly, IFT trains can be seen without ambiguity as electron dense particles found between the flagellar membrane and microtubule doublets (Video S1 and Fig. 2A, arrows on bottom panels). To determine doublet number, the fixed orientation of the axoneme relative to the PFR (Branche et al., 2006; Gadelha et al., 2006; Ralston et al., 2006) was exploited (Fig. 1A). Indeed, the presence of a thick projection that connects the B tubule of doublet 7 to the PFR makes doublet numbering straightforward (Fig. 1A). In case this projection was not visible, the proximal to distal orientation of the axoneme was determined by travelling through the volume knowing that dynein arms are orientated clockwise when starting from the base of the flagellum (Sherwin and Gull, 1989). Because several portions of flagella were encountered in a single volume, each axoneme was individually marked with a different colour and IFT trains were indicated in red (Fig. 2B,C)(Video S2). Most trains were found along doublets **3-4** and **7-8** and exhibited various lengths (Fig. 2B,C). We detected 52 trains closely associated to doublets **3-4** and 56 trains to doublets **7-8** in 27 distinct flagella (total cumulated axoneme length 166 µm, the average length of visible axoneme was 6.25 ± 4.21 µm). Only 2 trains were not found on these doublets (both were present on doublet 1). **All of the reconstructed axonemes from the FIB-SEM data were pooled in order to calculate an average train number in a typical axoneme. An average of 6.15 trains were found associated to doublets 3-4 and 6.6 trains on doublets 7-8.** This is below the expected numbers of 8.6 anterograde trains and 8.9 retrograde trains per flagellum deduced from their speed and frequency in live cells (Buisson et al., 2013), suggesting that some trains might be missed in the FIB-SEM analysis.

**Measurements of the length of IFT trains from the FIB-SEM data suggested the presence of two distinct populations on each set of doublets (Fig. 2D). This was confirmed by statistical analysis using R software with 61% of short trains (202 ± 94 nm) and 39% of longer trains of (822 ± 515 nm) associated to doublets 3-4 (n=52) whereas 68% of short trains (207 ± 99 nm) and 32% of longer trains of (968 ± 239 nm) were associated to doublets 7-8 (n=56) (Fig. 2D).**

**In most cases, the random orientation of flagella restricted the discrimination of individual doublets in the FIB-SEM volumes. However, we could find a few flagella whose positioning allowed the segmentation of individual doublets, hence offering the opportunity to see if IFT trains associate preferentially to doublet 3 or 4, or to doublet 7 or 8. One such example is presented at Figure 3 and corresponds to the cell in the centre of the field of view on Video S1. Doublets could be segmented over several microns and four trains were present along them. Trains annotated as IFT1, IFT2 and IFT4 were all found to be in closer proximity to doublet 7 compared to doublet 8 whereas train IFT3 was closer to doublet 4 than to doublet 3 (Fig. 3). This suggests that IFT trains could travel mostly on doublets 4 and 7. Unfortunately, this analysis was only feasible with very few flagella and cannot be taken as a generality.**

**Fig. 3.**
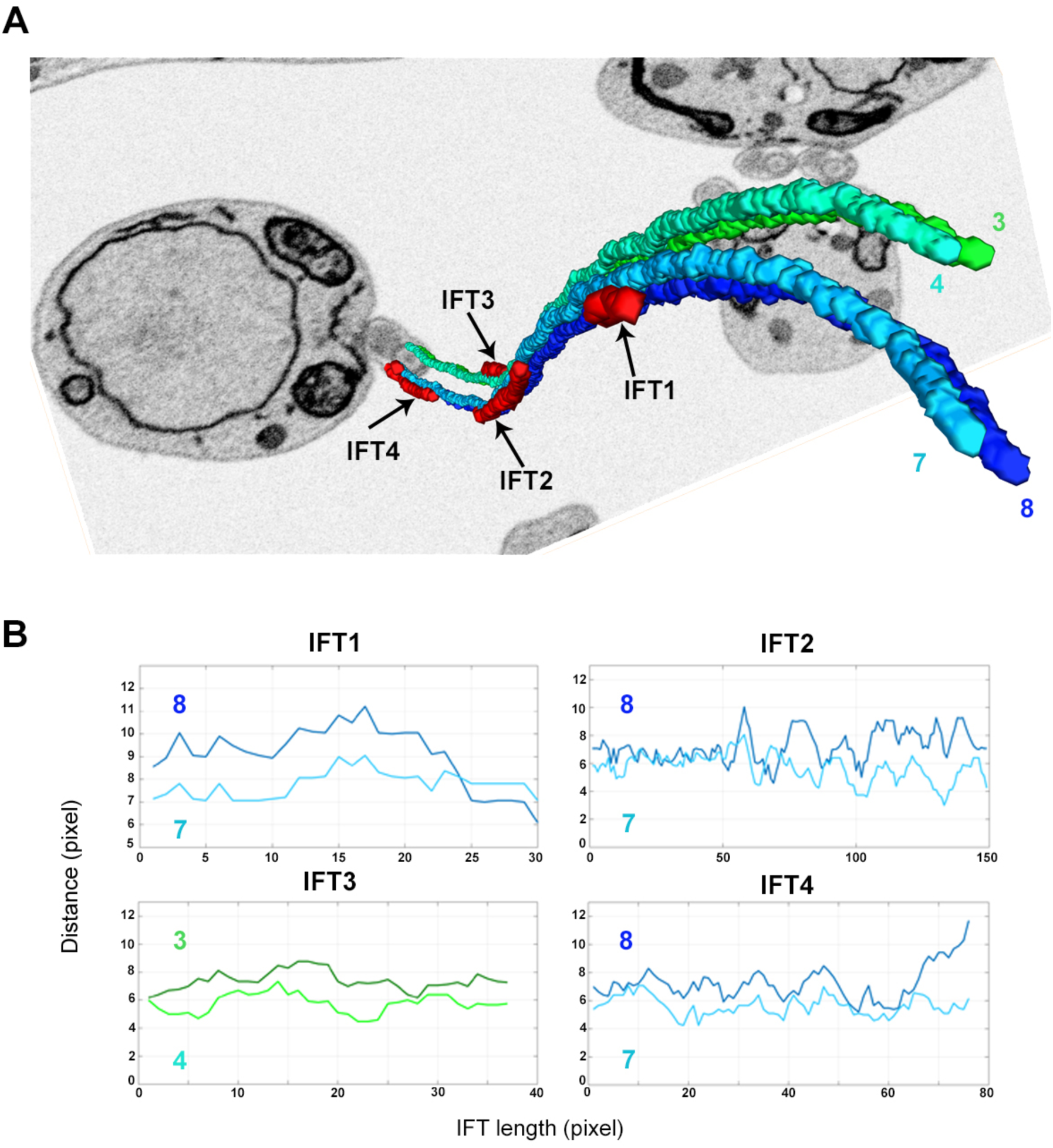
IFT trains are found closer to doublets 4 and 7 compared to doublets 3 and 8. (A) Three-dimensional view of a rare flagellum where individual doublets could be discriminated. Segmentation was performed to highlight the vicinity of the IFT trains (red) and the four microtubule doublets 3 (dark green), 4 (light green), 7 (light blue) and 8 (dark blue). (B) Plots representing the distances between the centre of the skeleton for the indicated IFT trains and that of the microtubules doublets along their length using the same colour code. In this flagellum, IFT trains are found closer to doublets 4 and 7.

**Overall, these results demonstrate that IFT trains are restricted to doublets 3-4 and 7-8 along the length of the flagellum in trypanosomes, with a possible preference for 4 and 7 at least in some flagella. The average train number and their average length are equivalent for the two sets of doublets, hence suggesting that similar trains travel on each of them, rather supporting the second hypothesis of anterograde and retrograde IFT trafficking taking place on each doublet. However, FIB-SEM does not give information on train direction and the possible presence of arrested trains (Stepanek and Pigino, 2016) cannot be ruled out and could interfere with the interpretation. Now that the spatial distribution of IFT trains has been clarified, we turned to live cell imaging in order to investigate their dynamics.**

### Immunofluorescence analysis indicates that IFT proteins are found on both sides of the axoneme

Measurements of flagellar sections by transmission electron microscopy showed that the doublets 4 and 7 are separated by 190 ± 11 nm (n=20), which is below the resolution of conventional light microscopy. To evaluate the feasibility of detecting IFT on these two separate **sides of the axoneme**, we tried different fixation protocols and examined the distribution of IFT proteins by immunofluorescence assay with antibodies against the IFT172 protein (Absalon et al., 2008) and the axonemal protein TbSAXO1 (Dacheux et al., 2012). Fixation of trypanosomes in paraformaldehyde followed by a post-fixation in methanol led to the detection of TbSAXO1 as a single thick line (second column, Fig. 4) as expected from immunogold analysis that showed that this protein was present throughout the axoneme (Dacheux et al., 2012). By contrast, IFT172 staining appeared as two parallel lines decorating both sides of the TbSAXO1 staining (third column, Fig. 4). This was observed in the single flagellum of G1 cells (Fig. 4A) and in both mature and growing flagella of duplicating cells (Fig. 4B). Methanol fixation results in dehydration and flattens the sample on the slide, possibly leading to a better separation of IFT trains allowing their detection as two separate lines by conventional light microscopy. This is promising because this suggests that IFT positioning on distinct doublets could be discriminated by light microscopy.

**Fig. 4.**
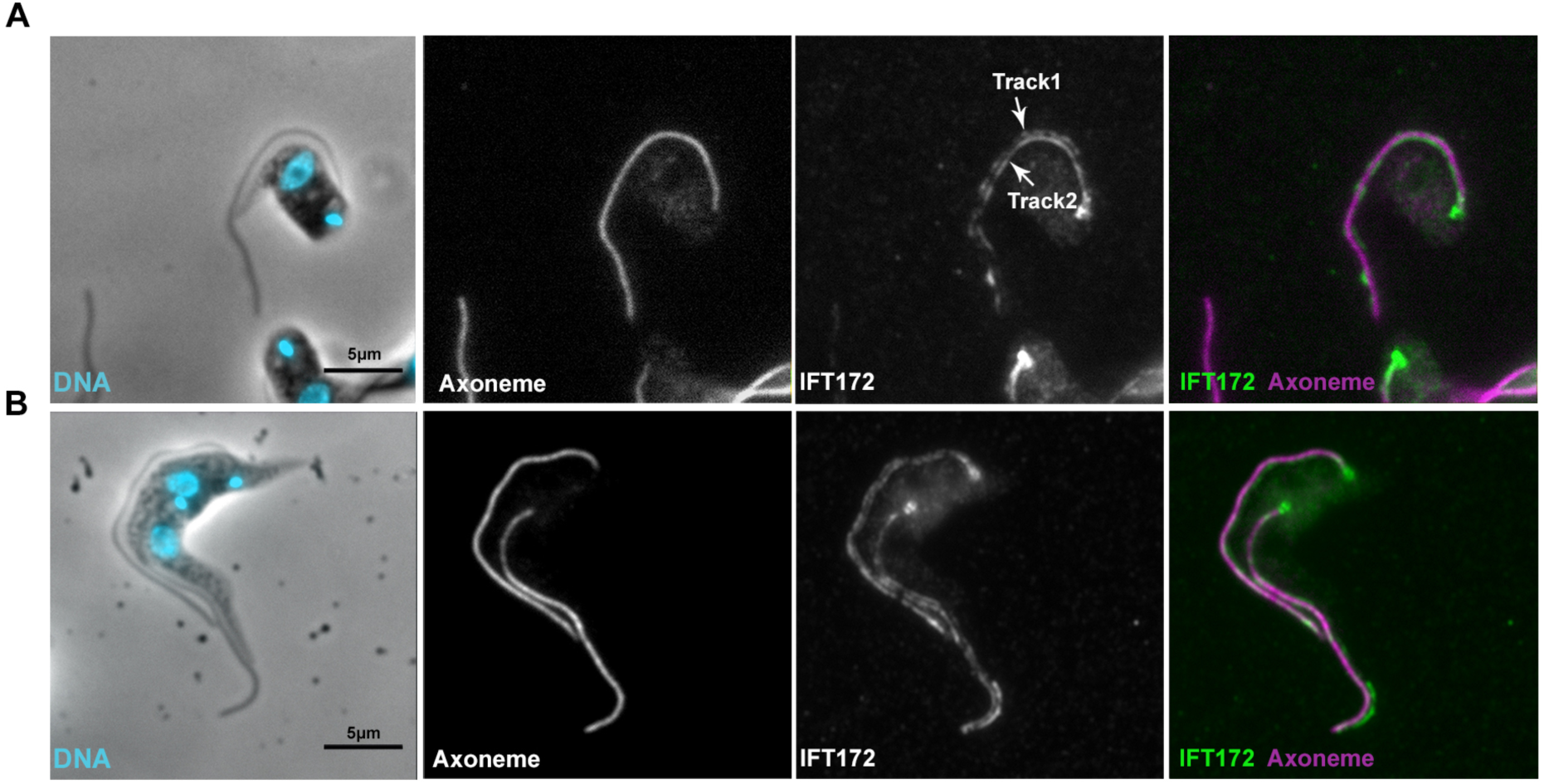
IFT proteins are found on two distinct lines along the axoneme in fixed cells. Control trypanosomes (strain expressing YFP::ODA8 (Bonnefoy et al., 2018)) were fixed in paraformaldehyde followed by methanol post-fixation and processed for immunofluorescence using a marker antibody for the axoneme (middle panel, magenta on the merged image) and a monoclonal antibody against IFT172 (right panel, green on the merged image). The first panel shows the phase contrast image merged with DAPI staining (cyan). **(A)** Cell with one flagellum. **(B)** Cell assembling a new flagellum. In both cases, a single continuous thick line is observed for the axoneme marker whereas discontinuous staining spreading on two close but distinct lines is visible for IFT172.

The next step was the investigation of IFT in live cells, which requires the expression of a fluorescent reporter in trypanosomes (Adhiambo et al., 2009; Bhogaraju et al., 2013; Buisson et al., 2013; Huet et al., 2014). Cell lines expressing GFP::IFT52 have been used reproducibly to detect IFT (Absalon et al., 2008; Buisson et al., 2013) but in this set-up, GFP::IFT52 was expressed from the ribosomal DNA locus with a strong promoter (Wirtz and Clayton, 1995). This led to bright flagellar signal but also to significant cytoplasmic signal (Buisson et al., 2013). To avoid any risk of potential artefacts due to over-expression, an *in situ* tagging approach (Dean et al., 2017; Kelly et al., 2007) was selected to generate a cell line expressing IFT81 fused to mNeonGreen (mNG)(Shaner et al., 2013) from its endogenous locus. IFT81 is a well-known member of the IFT-B complex involved in tubulin binding and transport (Bhogaraju et al., 2013; Kubo et al., 2016) and was shown previously to traffic within the trypanosome flagellum (Bhogaraju et al., 2013). Western blotting with an anti-mNG antibody demonstrated that the fusion protein displayed the expected mobility on SDS-PAGE (Fig. S1A). Live analysis showed the classic distribution with a strong signal at the base of the flagellum and train trafficking in both anterograde and retrograde direction within the flagellum that was detected by kymograph analysis (Video S3 & Fig. S1B-C). Kymograph analysis revealed that the frequency and speed of anterograde trains (Fig. S1D-E) was similar to what was reported previously for other IFT-B proteins (Bhogaraju et al., 2013; Buisson et al., 2013; Huet et al., 2014). Finally, applying the paraformaldehyde/methanol fixation protocol followed by direct observation of the mNG::IFT81 fluorescent signal led to the detection of two parallel lines within flagella (Fig. S1F). The behaviour of mNG::IFT81 is therefore comparable to that of GFP::IFT52 and both cell lines were used for imaging IFT in trypanosomes.

### High-resolution live cell imaging reveals bidirectional IFT on two sides of the axoneme

These results are encouraging because they suggest that live imaging by light microscopy could permit the visualisation of IFT on two **sides of the axoneme**. We turned to super-resolution based on structured illumination microscopy (SIM)(Gustafsson, 2000; Gustafsson et al., 2008) to visualise IFT trafficking. SIM imaging provided the spatial resolution and demonstrated the existence of two parallel lines within the flagellum for fluorescent mNG::IFT81 in live cells (Fig. S2A). Next, we tried to record IFT trafficking over time using the GFP::IFT52 cell line (Fig. S2B and Video S4). This demonstrated that IFT trains move on **two distinct parallel lines** in trypanosomes, in agreement with the electron microscopy data. However, the acquisition time (800 milliseconds) was not compatible with the rapid speed of IFT in trypanosomes (∼2 µm/s for anterograde and ∼5 µm/s for retrograde transport) and images of multiple trains overlapped, precluding analysis (Video S4).

To find an appropriate compromise between sufficient resolution and fast acquisition, “high-resolution” imaging was performed using objectives with superior numerical aperture (1.49 NA for GFP::IFT52, imaging performed at the Janelia Farm Research Campus, 1.57 NA for mNG::IFT81, imaging performed at the Institut Pasteur)(Li et al., 2015). In these conditions, the theoretical resolution for a green fluorescent molecule should be ∼160 nm, a value compatible with the discrimination of IFT trafficking on doublets separated by 190 nm. Remarkably, examination of cells expressing mNG::IFT81 (Video S5) or GFP::IFT52 (Video S6) with the high numerical aperture objectives revealed the presence of trains on two **parallel lines** within the flagellum. This was clearly confirmed with time projections (Fig. 5A and Fig. S3A). Closer examination of IFT trafficking demonstrated that anterograde and retrograde IFT trafficking was taking place on each of these **lines** (Fig. 5B, Fig. S3B, Video S5 and Video S6). This was further supported by kymograph analysis that showed the presence of distinct anterograde and retrograde trains on each **side of the axoneme** (Fig. 5C & Fig. S3C). These results demonstrate that IFT takes place on two separate tracks in the trypanosome flagellum presumably corresponding to doublets **3-4** and **7-8**, and that anterograde and retrograde trafficking occurs on each of them. **This supports the hypothesis of bidirectional IFT on both sets of doublets and probably invalidates the model with one track for each direction. However, we cannot formally exclude that anterograde trafficking takes place on 4 and retrograde on 3/or the opposite, and similarly with 7 and 8.**

**Fig. 5.**
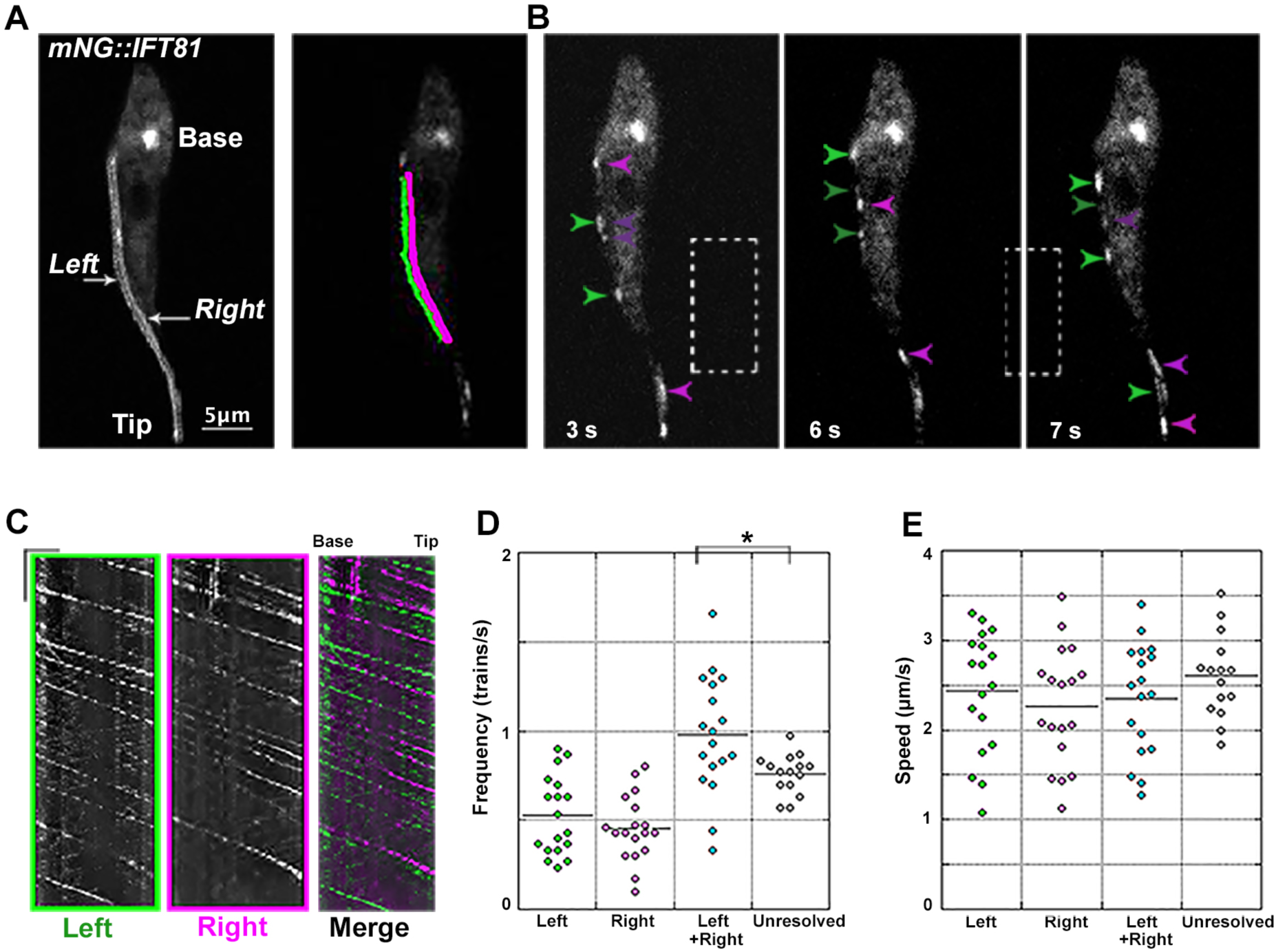
Bidirectional IFT trafficking takes place on two sides of the axoneme in live trypanosomes expressing mNG::IFT81. **(A)** Temporal projection of a stack of images corresponding to Video S5 showing the presence of two parallel lines for IFT in the flagellum in addition to the pool of IFT at the base. The left (green) and right (magenta) sides were defined after orientating the cell with the posterior end on top of the image and the flagellum on the left-hand side. **(B)** Still images from Video S5 of live trypanosomes expressing mNG::IFT81 imaged at high-resolution. Green and magenta arrowheads indicate trains on the left and right sides with light arrowheads pointing at anterograde trains and darker arrowheads showing retrograde trains. The time point for each image is indicated. **(C)** Kymograph analysis of the same cell showing trafficking on the left side (green), on the right side (magenta) and the merged images for the region of interest indicated in A. Scale bars are 2.5 µm for length (horizontal bar) and 2.5 seconds for time (vertical bar). **(D)** Dot plot of the frequency of anterograde IFT trains visible on the left (green) and the right (magenta) side, the sum of both (cyan) and from videos where only one track was visible (unresolved, grey). **(E)** Same representation but for the speed of anterograde trains. Only statistically significant differences are shown.

To be able to quantify and compare IFT train trafficking on each **side of the axoneme**, it was necessary to find a way to identify them. To do that, the cellular asymmetry of trypanosomes was exploited. Cells were orientated with the posterior end towards the top of the image and with the flagellum lying on the left-hand side, hence defining a left and a right **side** (Fig. 5A and Fig. S3A). Quantification of anterograde IFT train trafficking showed a similar frequency close to 0.5 anterograde train per second on the left and on the right **side** for both fluorescent proteins (Fig. 5D, Fig. S3D and Table 1). There was no statistically significant difference between these two parameters in cells expressing mNG::IFT81 (one-way ANOVA test, p=0.29) or GFP::IFT52 (p=0.14). The anterograde speed was ∼2.5 µm per second on each **side** in both cell lines (Fig. 5E, Fig. S3E and Table 1) although we noted a trend towards slower IFT speed by 10-15% on the right **side** (p=0.20 for mNG::IFT81 and p=0.067 for GFP::IFT52). Trains trafficking simultaneously on each **side** might not be discriminated with conventional light microscopy. Hence, the anterograde IFT train frequency was compared using videos of mNG::IFT81 cells acquired in the same experiment but where two **sides** could not be discriminated (Fig. 5E, “unresolved”). The frequency calculated from the total of left and right kymographs in cells where the two **sides** could be detected was consistently higher (0.98 anterograde trains per second) compared to cells where only one **line** was visible (0.76 anterograde trains per second)(Table 1) with statistical confidence (p=0.02). This result suggests that up to 15% of trains could be missed when IFT was imaged at low resolution. By contrast, their average train speed was indistinguishable (Table 1, p=0.21), suggesting that the “missed” trains are not a specific population at least in terms of speed.

**Table 1.**
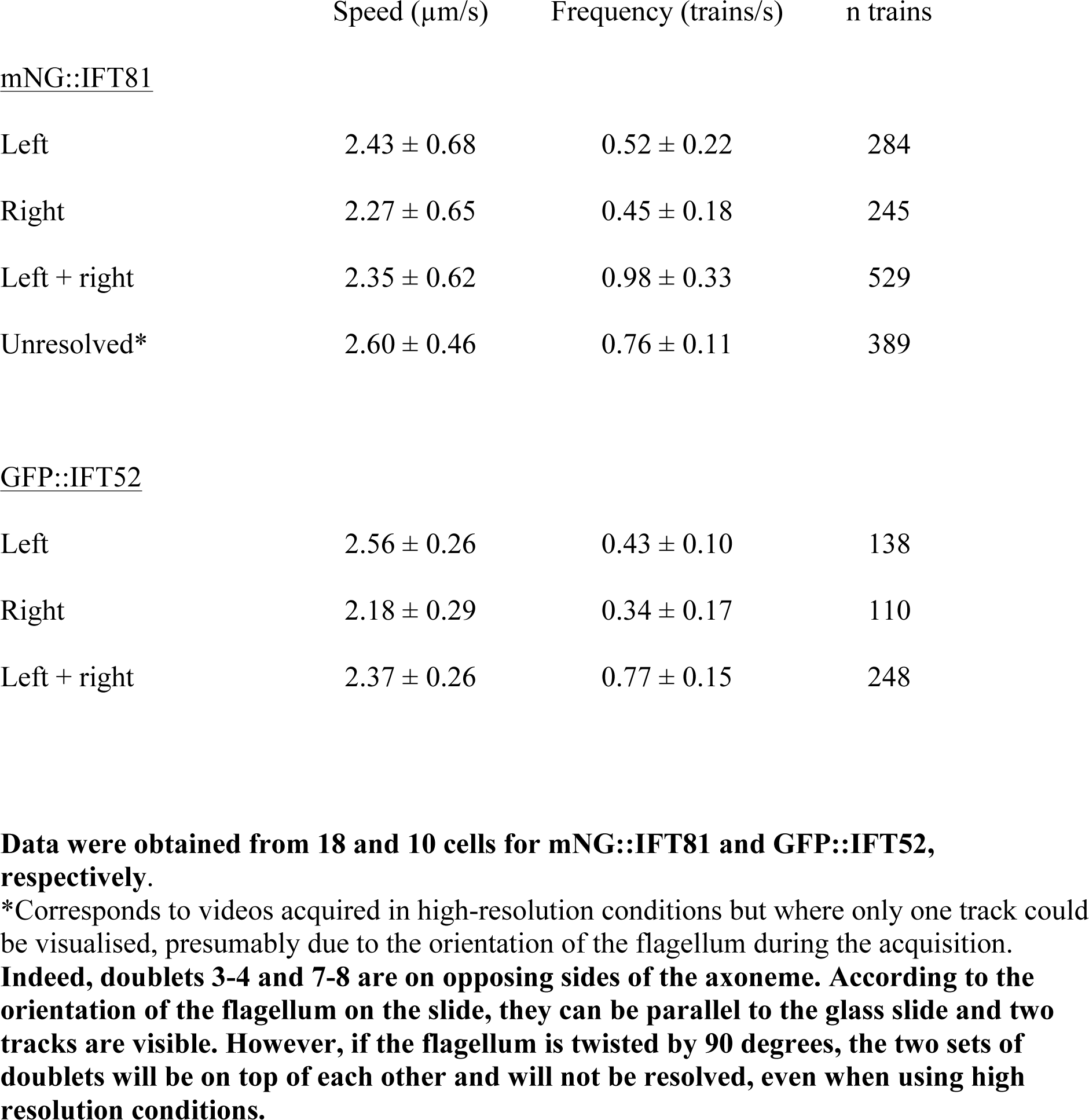
**Speed and frequency of anterograde IFT trafficking**

| | Speed ($\mu\text{m/s}$) | Frequency (trains/s) | n trains |
| --- | --- | --- | --- |
| <u>mNG::IFT81</u> |  |  |  |
| Left | $2.43 \pm 0.68$ | $0.52 \pm 0.22$ | 284 |
| Right | $2.27 \pm 0.65$ | $0.45 \pm 0.18$ | 245 |
| Left + right | $2.35 \pm 0.62$ | $0.98 \pm 0.33$ | 529 |
| Unresolved* | $2.60 \pm 0.46$ | $0.76 \pm 0.11$ | 389 |
| <u>GFP::IFT52</u> |  |  |  |
| Left | $2.56 \pm 0.26$ | $0.43 \pm 0.10$ | 138 |
| Right | $2.18 \pm 0.29$ | $0.34 \pm 0.17$ | 110 |
| Left + right | $2.37 \pm 0.26$ | $0.77 \pm 0.15$ | 248 |

**Data were obtained from 18 and 10 cells for mNG::IFT81 and GFP::IFT52, respectively.**
\*Corresponds to videos acquired in high-resolution conditions but where only one track could be visualised, presumably due to the orientation of the flagellum during the acquisition.
**Indeed, doublets 3-4 and 7-8 are on opposing sides of the axoneme. According to the orientation of the flagellum on the slide, they can be parallel to the glass slide and two tracks are visible. However, if the flagellum is twisted by 90 degrees, the two sets of doublets will be on top of each other and will not be resolved, even when using high resolution conditions.**

Although retrograde transport was detected in almost all videos (Video S5 and Video S6), the lower intensity of these trains made their quantification quite challenging, especially for the frequency. Nevertheless, it was possible to estimate the speed of the brightest retrograde trains, which was between 4 and 5 µm per second, in agreement with data obtained using conventional imaging (Buisson et al., 2013).

**Although our live cell imaging data reveals two separate paths for IFT, we wanted to confirm that the IFT motors that connect IFT particles to microtubules exhibit the same. Molecular motors of the kinesin and dynein family transport IFT proteins in the anterograde and retrograde directions, respectively. We investigated the movement of the dynein motor and monitored its association to the IFT proteins. First, we monitored the movement of dynein using the heavy chain 2.2 (DHC2.2) fused to GFP (Blisnick et al., 2014) using high-resolution imaging. As for IFT52 and IFT81, trafficking on two parallel lines on each side of the axoneme was detected (Fig. 6A & Video S7, n=14). Kymograph analysis confirmed independent movement in anterograde and retrograde direction on both sides (Fig. 6B). We next set up dual-colour imaging to visualise simultaneously mNG::IFT81 and the dynein heavy chain 2.1 (DHC2.1) tagged *in situ* with TandemTomato (TdT) in live cells. Overall, both signals seem to co-localise in moving and standing trains (Fig. 6C & Video S8). Analysis of 10 cells revealed that 93% and 81% of the anterograde traces were positive for both mNG::IFT81 and TdT::DHC2.1 in the anterograde (n=273 trains) and retrograde (n=41) direction. Lower values for retrograde trains probably reflects the weaker fluorescent signal produced by the TdT::DHC2.1 compared to mNG::IFT81. Unfortunately, the same experiment could not be done with IFT kinesins since tagging disrupted protein localisation (data not shown).**

**Fig. 6.**
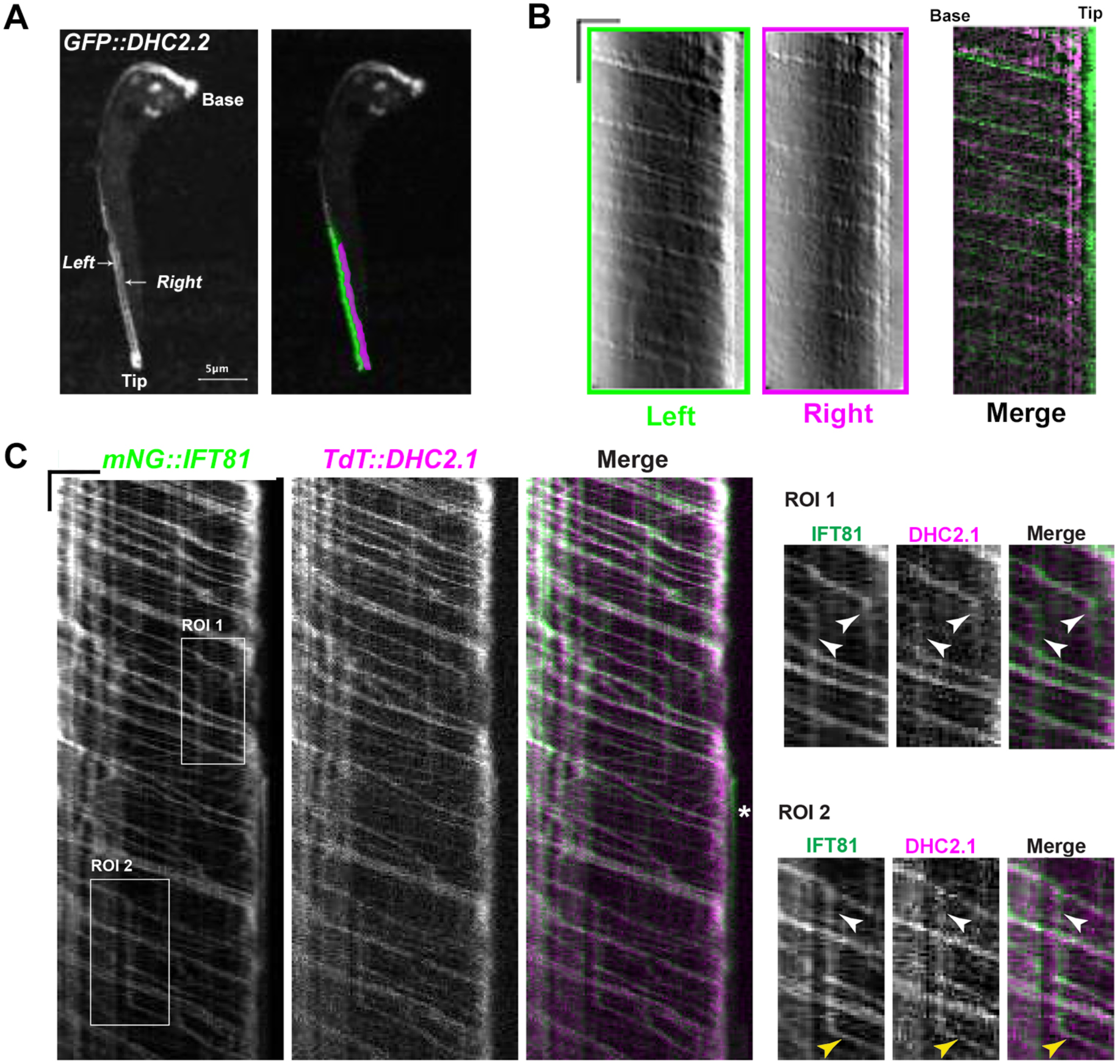
The IFT dynein travels on both sides of the axoneme and is associated to IFT proteins. (A) Temporal projection of the flagellum in a cell expressing GFP::DHC2.2. The region of interest at the tip of the flagellum is indicated, with left (green) and right (magenta) sides highlighted. (B) The kymographs are shown for the left and right side in the corresponding colour. Scale bars are 2.5 µm for length (horizontal bar) and 5 seconds for time (vertical bar). (C) Kymograph analysis of a cell expressing mNG::IFT81 (left panel) and TdT::DHC2.1 (centre panel). On the merged panels mNG::IFT81 is shown in green and TdT::DHC2.1 in magentas. Both proteins co-localise in moving but also in standing trains shown in the regions of interest (ROI). White arrowheads indicate arrested trains while the yellow arrowheads point at an arrested train that started moving again. Even in these conditions, mNG::IFT81 and TdT::DHC2.1 remain associated. The white star on the merged panel indicates a rare example of mNG::IFT81 material that is not associated with TdT::DHC2.1 and that remains immotile at the far distal end of the flagellum.

### Conversion of anterograde to retrograde train takes place on the same side of the axoneme

To visualise the conversion of anterograde to retrograde transport, we looked at the distal end of the flagellum of cells expressing mNG::IFT81 (Fig. S4A & Video S9). On both left and right **sides**, the arrival of large anterograde trains at the distal end of the flagellum was clearly visible. This was followed by a lag phase (seen as vertical lines on the kymograph) where the fluorescent material remained at the distal end but its intensity progressively went down while multiple retrograde trains were released during a ∼3-4 second period (Fig. S4B). It should be noted that the anterograde trains do not all stop exactly at the same place (merged panel, Fig. S4B). Data suggested that anterograde trains convert to retrograde trains on the same track and that no exchange of IFT-B material could be detected between left and right sides of the axoneme. **However, the high frequency of IFT in both directions and the relatively weak signal of retrograde trains make it difficult to reach a firm conclusion. We therefore used the GFP::IFT52 expressing cell line where the signal is brighter, hence facilitating the detection of retrograde transport. The distal end of the flagellum was bleached and IFT was recorded in this portion (Fig. 7 & Video S10). This led to a significant improvement of the signal-to-noise ratio and helped the visualisation of retrograde trains on videos and on kymographs. The arrival of several anterograde trains and their conversion to retrograde trains was tracked carefully examining video images and performing kymograph analysis (Fig. 7 & Video S10). We could not see an exchange of sides scrutinizing 12 movies acquired in these conditions, following dozens of trains in each.**

**Fig. 7.**
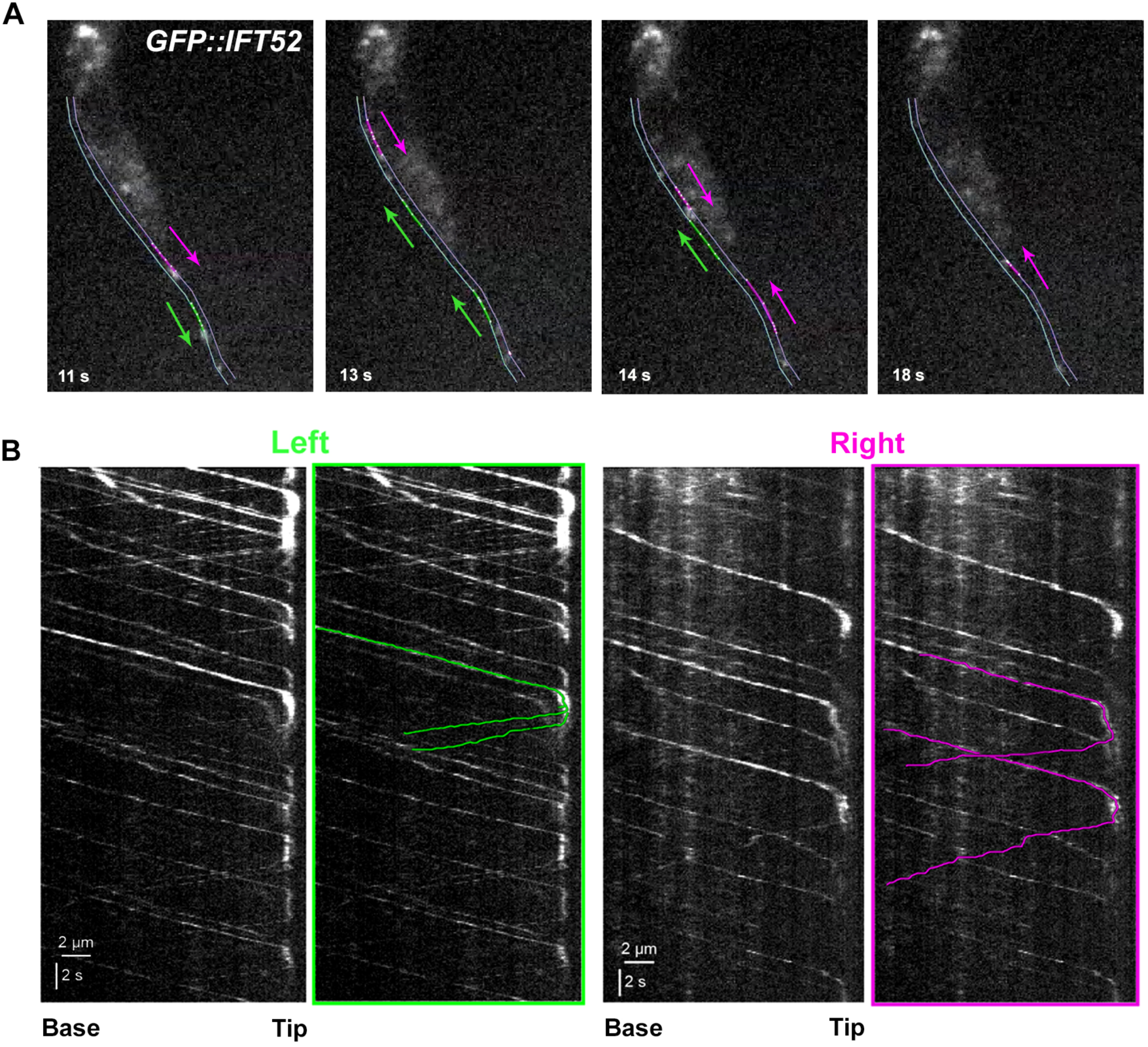
Anterograde trains are converted to retrograde trains whilst remaining on the same track. IFT trafficking was analysed in the GFP::IFT52 expressing cell line. The tip of the flagellum was bleached and the arrival of new anterograde trains was monitored (see corresponding Video S10). For technical reasons, the bleaching event could not be recorded and only the recovery phase can be presented. (A) The panel shows still images corresponding to the indicated times from Video S10. The thin green and magenta lines indicate the left and right side of the axoneme, respectively. Image 11s: Two anterograde trains are marked with a green (left side) or a magenta (right side) line and the corresponding arrows indicate their direction. Image 13s: The left anterograde train converted to two retrograde ones that remain on the same left side of the flagellum. A second anterograde train arrives on the right side. Image 14s: The first left retrograde train has gone out of the field of view and the second one also progresses towards the base of the flagellum. The first anterograde train of the right side has converted to a retrograde one that also remains on the right side. Image 18s: All retrograde trains on the left side are not in the field of view anymore, as well as the first one on the right side. A new retrograde train is present on the right side issued from the second anterograde train. Once again, conversion took place on the same side of the axoneme. (B) Kymograph analysis for the indicated left and right side of the flagellum. The original kymograph is shown on the left of each panel and the three annotated trains described above are marked in colour.

Overall, these results obtained with two different cell lines (expression of mNG::IFT81 and GFP::IFT52) at two different imaging centres (Janelia Farm, USA and Institut Pasteur, France) using different techniques demonstrate that IFT takes place on two distinct **sides of the flagellum** and that anterograde and retrograde trafficking occur on each of them.

## DISCUSSION

*T. brucei* is only the second organism where IFT trains have been visualised in both light and electron microscopy. Here we reveal train distribution on doublets **3-4 and 7-8** using FIB-SEM **(with possibly a preference for 4 and 7)** and demonstrate bidirectional trafficking on two **separate sides of the axoneme** by high-resolution live cell imaging. These two sides presumably correspond to doublets 3-4 and 7-8 but definitive confirmation will require the identification of specific markers of each of these doublets at the axoneme level.

FIB-SEM detected two distinct populations of trains on each set of doublets. The first possibility is that long trains correspond to anterograde ones and short trains to retrograde ones, as expected from the distinctive size on videos and kymographs (Buisson et al., 2013). The higher frequency of short trains (2.3 to 1.5-fold) is consistent with the higher abundance of retrograde trains detected during live imaging (Buisson et al., 2013). However, the total number of IFT trains seen in FIB-SEM (12.5 per flagellum) is lower compared to what was detected by live imaging (17.5). **Moreover, the average length of long trains is close to 1 µm and this is highly unusual for anterograde trains seen on kymographs, in contrast to arrested trains that look larger (Figure S4).** Intriguingly, two populations of IFT trains were also found using electron tomography analysis of *Chlamydomonas* flagella, with average lengths fairly close to what is reported here for trypanosomes (Pigino et al., 2009). Long trains were initially thought to correspond to anterograde trains because they accumulate in the *fla14* retrograde transport mutant where small trains disappear. However, correlative light and electron microscopy revealed that long trains correspond to standing material and that short particles correspond to anterograde trains (Stepanek and Pigino, 2016). Based on this, we propose that short and long trains observed by FIB-SEM in *T. brucei* flagella **could** correspond to anterograde trains and to standing material, respectively. A total of 7.2 short trains is found per flagellum, which is close to the predicted number of 8.6 (Buisson et al., 2013). Observation of kymographs indicated the presence of standing trains that look larger than moving trains (Fig. S4B, stars). It should be noted that this material does not remain stuck forever as it appears to be picked up by other anterograde trains after a few seconds (stars on Fig. S4B and Fig. S1E). In this scenario, retrograde trains would be missed either because they are too short or because their morphology is different and difficult to identify by FIB-SEM. In *Chlamydomonas*, retrograde trains appear less condensed and less regular compared to anterograde trains, and had been missed in conventional transmission electron microscopy until their identification by correlative techniques (Stepanek and Pigino, 2016).

The use of superior numerical aperture objectives appears to be an optimal compromise between speed of acquisition and spatial resolution. It revealed the existence of bidirectional IFT on two sides of the axoneme but it also showed that the frequency of IFT measured with conventional microscopy might be slightly underestimated. Moreover, it revealed the transition of anterograde trains to retrograde trains with IFT-B proteins spending up to 3-4 seconds at the distal end while being progressively associated to the emergence of several retrograde trains. This value is in agreement with previous low-resolution analysis based on photobleaching experiments (Buisson et al., 2013). In *Chlamydomonas* (Chien et al., 2017) and in *C. elegans* (Mijalkovic et al., 2017), IFT-B proteins also spend a few seconds at the distal end of microtubules before returning towards the base as components of several retrograde trains.

The restriction of IFT to a limited set of doublets is different from the situation encountered in *Chlamydomonas* where trains appear to be present on the majority of the doublets (Stepanek and Pigino, 2016). So far, IFT trains have only been visualised by electron microscopy in these two organisms. To understand the significance of IFT presence on all or only some doublets, it will be essential to determine which doublets are being used in different types of cilia in various organisms. This raises the question of why restricting IFT to some doublets. We propose a scenario where the restriction of IFT to some doublets would represent an evolutionary advantage by liberating the other doublets from constraints imposed by IFT train presence, thereby offering the opportunity for acquiring new structures or functions. In trypanosomes and related protists, the PFR is tightly associated to the axoneme (Hughes et al., 2012; Koyfman et al., 2011) and brings an essential contribution to flagellum motility (Bastin et al., 1998; Santrich et al., 1997). Restricting IFT to some doublets could allow the passage of other molecular motors on the remaining doublets. These could transport different cargoes, hence increasing the range of functions performed by cilia and flagella. In trypanosomes, an unusually large number of genes encoding for kinesins has been identified (Berriman et al., 2005) and several of the protein products have been localised to the flagellum (Chan and Ersfeld, 2010; Demonchy et al., 2009) or found in proteomic analyses of purified flagella (Broadhead et al., 2006; Oberholzer et al., 2011; Subota et al., 2014).

The trypanosome flagellum is attached to the cell body for most of its length towards the PFR side. Doublets 1-2-3-8-9 are towards the surface of the flagellum and the absence of IFT could favour interactions with host tissues. For example, parasites interact with the epithelium of the salivary glands of the tsetse fly via their flagellum and the development of electron-dense material resembling hemi-desmosomes (Tetley and Vickerman, 1985). In other organisms, the cilia of sensory neurons of *C. elegans* spring to mind. They are composed of a middle segment made of 9 doublet microtubules and of a distal segment with only singlet microtubules. Bidirectional IFT was reported on both segments without collisions (Snow et al., 2004). If some microtubules were used only for anterograde IFT and others only for retrograde IFT, it would provide a way to avoid collisions (Kuhns and Blacque, 2016).

Here, we show a striking functional difference between doublets 3-4 & 7-8 and the others that cannot sustain IFT. Although microtubule doublets look similar, discrete molecular and structural differences have been noted between them in several organisms (Heuser et al., 2012; Lin et al., 2012). This is also the case in trypanosomes where doublets 4, 5, 6 and 7 are physically linked to the PFR by different structures (Hughes et al., 2012; Sherwin and Gull, 1989) that could contain unique proteins (Imboden et al., 1995). Other molecular differences between doublets start to be unveiled with the recent example of CFAP43 and CFAP44, two proteins required for motility that have been located to doublets 5 and 6 using super-resolution microscopy (Coutton et al., 2018).

What could make doublets 4 and 7 different from the others and why would they be used for IFT? One possibility is that they contain biochemical information that is preferentially recognised by the IFT molecular motors. Promising candidates are post-translational modifications of tubulin such as (poly)glycylation or (poly)glutamylation. These are found at the surface of the tubulin dimer where one would expect interactions with molecular motors (Konno et al., 2012; Sirajuddin et al., 2014). Insights into a potential molecular mechanism are provided by *in vitro* experiments using engineered tubulin with various post-translational modifications. This revealed that recruitment and processivity of the IFT kinesin motor KIF17 was stimulated by polyglutamylation (Sirajuddin et al., 2014). Polyglutamylation and polyglycylation are overwhelmingly represented in cilia and flagella and their alteration affect these organelles (Bosch Grau et al., 2013; Lee et al., 2012; Pathak et al., 2011; Rogowski et al., 2009; Wloga et al., 2009). Mass spectrometry showed that trypanosome tubulin is extensively polyglutamylated, with variable numbers of glutamate residues added to both cytoplasmic and flagellar microtubules (Schneider et al., 1997). Investigating the role of tubulin glutamylation in the definition of microtubule heterogeneity will be an exciting but challenging future axis for research given the large number of enzymes involved in this process (Janke et al., 2005; van Dijk et al., 2007) including in *T. brucei* (Casanova et al., 2014).

## Methods

### Trypanosome cell lines and cultures

Cell lines used for this work were derivatives of *T. brucei* strain 427 cultured at 27°C in SDM79 medium supplemented with hemin and 10% foetal calf serum (Brun and Schonenberger, 1979). Cell line expressing GFP::DHC2.1 or GFP::DHC2.2 have been described previously (Blisnick et al., 2014). The cell line expressing GFP::IFT52 from the pHD430 vector (Absalon et al., 2008) under the control of the tet-repressor (produced by plasmid pHD360 (Wirtz and Clayton, 1995)) was transformed to express a Tandem Tomato::IFT81 fusion produced from its endogenous locus (Bhogaraju et al., 2013). For the generation of the mNeonGreen::IFT81 expressing cell line, the first 497 nucleotides of *IFT81* (Gene DB number Tb927.10.2640, without the ATG) were chemically synthesised (GeneCust, Luxembourg) and cloned in frame with the *mNeonGreen* gene (Shaner et al., 2013) within the HindIII and ApaI sites of p2675 vector (Kelly et al., 2007). The construct was linearised within the *IFT81* sequence with the enzyme XcmI and nucleofected (Burkard et al., 2007) in the wild-type 427 cell line, leading to an integration by homologous recombination in the endogenous locus and to expression of the full-length coding sequence of IFT81 fused to mNeonGreen. The same procedure was used for endogenous tagging of the heavy chain DHC2.1 subunit of the dynein motor using the first 497 bp of the coding sequence (Gene DB number Tb927.4.560 without the ATG) cloned in the p2845 vector containing the *TandemTomato* gene and linearized with Csi1. Transfectants were grown in media with the appropriate antibiotic concentration and clonal populations were obtained by limited dilution.

### Focused Ion Beam-Scanning Electron Microscopy (FIB-SEM)

Trypanosomes were fixed directly in medium with 2.5% glutaraldehyde (Sigma), cells were spun down, the supernatant was discarded and the pellet was incubated for 15 minutes in fixation buffer made of 2.5% glutaraldehyde and 4% paraformaldehyde in cacodylate 0.1M buffer (pH 7.4). Samples were washed 3 times with 0.1M cacodylate buffer (5 minutes each) and post fixed with 1% osmium (EMS) and 1.5% potassium ferrocyanide (Sigma) in 0.1M cacodylate buffer for 1h. Samples were treated for 30 minutes with 1% tannic acid (Sigma) and 1h with 1% osmium tetroxide (EMS), rinsed in water and dehydrated in ethanol (Sigma) series of 25%, 50%, 75%, 90% and 100% (15 minutes each). Cells were embedded in epoxy resin (EMS) after 48h at 60°C of polymerization. Embedded samples were mounted on aluminium stubs. Blocks were trimmed with glass knives in such a way that exposure of vertical faces allowed lateral milling by Focused Ion Beam FIB. Tomographic datasets were obtained using a FESEM Zeiss Auriga microscope equipped by a CrossBeam workstation (Carl Zeiss) and acquired using ATLAS 3D software (Carl Zeiss). For milling with the focused Ga-ion beam, the conditions were as follows: 0.5–1nA milling current of the Ga-emitter, leading to the removal of 10 nm at a time from the epoxy resin. SEM images were recorded with an aperture of 60 µm in the high-current mode at 1.5 or 2 kV of the in-lens EsB detector with the EsB grid set to −1000 V. Depending on the respective magnification, voxel size was in a range between 10 and 20 nm in *x*/*y* and 10 nm in *z*. Contrast of the images was inverted to conventional bright field. Two different persons performed the manual annotation of IFT trains. These were defined as electron dense structures sandwiched between the axoneme and the flagellum membrane and present on a minimum of 3 consecutive slices (30 nm). Densities associated to membrane distortions were excluded. Trains were defined as different when separated by a minimum of 3 slices (30 nm).

### Data processing and 3-D-reconstruction

Alignment of image stacks was done with the open source software ImageJ for data alignment (Schneider et al., 2012) and Amira Software for visualization (FEI Thermofisher, v6.0.1). Segmentation and 3-D reconstructions were performed semi-automatically using Amira software and were corrected manually.

### Length measurement of IFT trains

Segmentations of flagella and IFT trains were first split according to their segmented colours then skeletonized in ImageJ (Schneider et al., 2012) using the Skeletonize 3D plugin (Arganda-Carreras et al., 2010). The number of voxels composing the generated skeletons was computed in ImageJ using the Object Counter 3D plugin (Bolte and Cordelieres, 2006). The length of analysed biological structures (flagella and IFT trains) was calculated as the number of voxels constituting the skeletonized structure multiplied by the size of the voxel.

### Distance measurement between IFT trains and individual doublets

IFT trains and doublets 3, 4, 7 and 8 were manually segmented on one flagellum to assess the distance between the IFT trains and the doublets. IFT train and doublet segmentations were skeletonized in ImageJ (Schneider et al., 2012) using the Skeletonize 3D plugin (Arganda-Carreras et al., 2010). We then computed in 3D the minimum distance between each voxel of the IFT train skeletons and the voxels of the doublet skeletons. This allowed to plot along the IFT train its distance with the various doublets in order to determine the closest one.

### Statistical analyses

In absence of other indications, all errors correspond to the standard deviation of the population. Anova tests were performed using the appropriate tool in Kaleidagraph v4.5.2. Populations of IFT trains on doublets 4 and 7 were analysed separately with the statistical analysis software R (Team, 2014) using the normalMixEM algorithm of the Mixtools package (version 1.1.0)(Benaglia et al., 2009) to check whether they were composed of sub-populations or not. This algorithm based on expectation maximisation estimates the mean and standard deviation values of Gaussian sub-populations and eventually converges to a solution if such sub-populations exist. Convergence was reached in 27 and 13 iterations for IFT trains on doublet 4 and 7 respectively.

### Structured illumination microscopy (SIM)

Trypanosomes expressing the mNG::IFT81 fusion protein were spread on glass coverslips in medium and SIM was performed on a Zeiss LSM780 Elyra PS1 microscope (Carl Zeiss, Germany) using 100×/1.46 oil Plan Apo objective and an EMCCD Andor Ixon 887 1 K camera for the detection at the Institut Pasteur. Fifteen images per plane per channel (five phases, three angles) were acquired to perform the SIM image. SIM image was processed with ZEN software. The SIMcheck plugin (Ball et al., 2015) in ImageJ (Schneider et al., 2012) was used to evaluate the acquisition and the processing parameters.

### High-resolution imaging of IFT trafficking

The cell line expressing GFP::IFT52 and Tandem Tomato::IFT81 was grown in standard conditions and samples were mounted between glass and coverslip for observation on a custom built microscope (Gustafsson, 2000; Gustafsson et al., 2008) based on a Zeiss AxioObserver D1 stand equipped with an UAPON100XOTIRF 1.49 NA objective (Olympus) and an Orca Flash 4.0 sCMOS camera (Hamamatsu). GFP fluorophores were excited with a 488 nm laser (500 mW, SAPPHIRE 488-500, Coherent) and detected through an adequate emission filter (BP 500-550 nm). The sequence contains a series of 300 images exposed for 20 milliseconds each, for a total duration of 17.7 s. Kymographs of individual paths of IFT were extracted using Fiji (Schindelin et al., 2012). The two IFT tracks were manually annotated as segmented lines on the temporal maximal intensity projection of the sequence. These two lines were then used to re-slice the sequence data, generating the kymographs that were analysed using Icy (de Chaumont et al., 2012). Cell lines expressing mNG::IFT81 or GFP::DHC2.2 were grown in standard conditions and samples were mounted between glass and quartz coverslip (Cover glasses HI, Carl Zeiss, 1787-996). For movie acquisition, a spinning disk confocal microscope (UltraVIEW VOX, Perkin-Elmer) equipped with an oil immersion objective Plan-Apochromat 100x/1.57 Oil-HI DIC (Carl Zeiss) was used. Movies were acquired using Volocity software with an EMCCD camera (C-9100, Hamamatsu) operating in streaming mode. The samples were kept at 27°C using a temperature controlled chamber. Sequences of 30 seconds were acquired with an exposure time of 100 milliseconds per frame. Kymographs were extracted and analysed with Icy software (de Chaumont et al., 2012) using the plug-in Kymograph Tracker 2 (Chenouard et al., 2010). The cells were positioned in the same orientation with the posterior end on top and the flagellum on the left-hand side to define the left and right sides. The two traces left by IFT trains were manually defined as Region of Interest using the temporal projection.

Dual colour acquisitions were performed with the cell line expressing mNG::IFT81 and TdT::DHC2.1on a spinning disk fluorescence microscope (UltraVIEW VOX, Perkin-Elmer) equipped with confocal scanning head (CSU X1, Yokagawa Electrics) and a 100x 1.4NA objective controlled by the Volocity software. The CSU X1 incorporates a dichroic mirror that sends emitted light on two EM-CCD cameras (ImagEM C9100, Hamamatsu), separately for wavelength below and above 550nm. This allows for simultaneous dual colour imaging for red and green channels.

### Immunofluorescence imaging

For paraformaldehyde-methanol fixation, cultured parasites were washed twice in SDM79 medium without serum and spread directly onto poly-L-lysine coated slides (Fisher Scientific J2800AMMZ). Cells were left for 10 minutes to settle prior to treatment with 1 volume 4% PFA solution in PBS at pH 7. After 5 minutes, slides were washed briefly in PBS before being fixed in pure methanol at a temperature of −20°C for 5 minutes followed by a rehydration step in PBS for 15 minutes. For immunodetection, slides were incubated for 1 h at 37°C with the appropriate dilution of the first antibody in 0.1% BSA in PBS; mAb25 recognises the axonemal protein TbSAXO1 (Dacheux et al., 2012; Pradel et al., 2006) and a monoclonal antibody against the IFT-B protein IFT172 (Absalon et al., 2008). After 3 consecutive 5-minute washes in PBS, species and subclass-specific secondary antibodies coupled to the appropriate fluorochrome (Alexa 488, Cy3, Jackson ImmunoResearch) were diluted 1/400 in PBS containing 0.1% BSA and were applied for 1 h at 37°C. After washing in PBS as indicated above, cells were stained with a 1µg/ml solution of the DNA-dye DAPI (Roche) and mounted with the Slowfade antifade reagent (Invitrogen). Slides were immediately observed with a DMI4000 microscope (Leica) with a 100X objective (NA 1.4) using a Hamamatsu ORCA-03G camera with an EL6000 (Leica) as light excitation source. Image acquisition was performed using Micro-manager software and images were analysed using ImageJ (National Institutes of Health, Bethesda, MD).

### Western blot

Cells were washed once in Phosphate Buffer Saline (PBS). Laemmli loading buffer was added to the cells and samples were boiled for 5 minutes. 20µg of protein were loaded into each lane of a Criterion™ XT Bis-Tris Precast Gel 4-12% (Bio-Rad, UK) for SDS-Page separation. XT-Mops (1X) diluted in deionised water was used as a running buffer. Proteins were transferred onto nitrocellulose membranes using the BioRad ^®^Trans-Blot Turbo™ blotting system (25V over 7 minutes). The membrane was blocked with 5% skimmed milk for one hour and then incubated with the monoclonal anti-mNeonGreen (32F6) primary antibody (ChromoTek, Germany) diluted 1/1000 in 0.05% PBS-Tween (PBST). As a loading control the anti-PFR L13D6 monoclonal antibody (Kohl et al., 1999) diluted 1/25 was used. After primary antibody incubation, three washes of 5 minutes each were performed in 0.05% PBST followed by secondary antibody incubation. Anti-mouse secondary antibody coupled to horseradish peroxidase, diluted to 1/20,000 in 0.05% PBST containing 0.1% milk, and the membrane was incubated with this for 1 hour. The Amersham ECL Western Blotting Detection Reagent Kit (GE Healthcare Life Sciences, UK) was used for final detection of proteins on the membrane.

## Acknowledgements

We thank Sylvie Perrot (Institut Pasteur Paris) for the preparation of samples for FIB-SEM, Audrey Salles (Institut Pasteur Paris) for the SIM acquisition performed on the Zeiss Elyra SP1 system, Derrick Robinson (Bordeaux University, France) for providing the Mab25 antibody and Linda Kohl (Museum of Natural History, Paris) for critical reading of the manuscript. We thank Adrien Vuilaume (Carl Zeiss, France) for support and interest for this project. We are grateful to the Photonic Bioimaging and Ultrastructural Bioimaging facilities for access to their equipment. Some SIM and high-resolution microscopy work was performed at the Advanced Imaging Centre, Janelia Research Campus, jointly sponsored by the Howard Hughes Medical Institute and the Gordon & Betty Moore Foundation. We are grateful to Lin Shao for training and advice. The generosity and the reactivity of Jim Morris (Clemson University, USA) are warmly acknowledged for having made this project feasible. E.B. is supported by fellowships from French National Ministry for Research and Technology (doctoral school CDV515) and from La Fondation pour la Recherche Médicale (FDT20170436836). C. F. was supported by fellowships from French National Ministry for Research and Technology (doctoral school CDV515) and from La Fondation pour la Recherche Médicale (FDT20150532023). This work is funded by ANR grants (11-BSV8-016 and 14-CE35-0009-01), by La Fondation pour la Recherche Médicale (Equipe FRM DEQ20150734356) and by a French Government Investissement d’Avenir programme, Laboratoire d’Excellence “Integrative Biology of Emerging Infectious Diseases” (ANR-10-LABX-62-IBEID). We are also grateful for support for FESEM Zeiss Auriga and Elyra PS1 equipment from the French Government Programme Investissements d’Avenir France BioImaging (FBI, N° ANR-10-INSB-04-01) and from a DIM-Malinf grant from the Région Ile-de-France. Travel to the AIC, Janelia Farm Research Campus was supported by the Citech (Centre d’Innovation et Recherche Technologique) of the Institut Pasteur.

## Author contributions

E. Bertiaux segmented FIB-SEM data, conducted live cell acquisition with the mNG::IFT81 or mNG::IFT81/TdT::DHC2.1 cell lines, extracted and analysed the kymographs and performed the statistical analyses on IFT trafficking and contributed to figure preparation. A. Mallet acquired the FIB-SEM data, segmented them and produced the corresponding videos as well as the analysis of kymographs and the acquisition of GFP::DHC2.2 images at high resolution. C. Fort prepared samples for FIB-SEM and carried out several segmentations, conducted the GFP::IFT52 acquisition and performed kymograph analyses. T. Blisnick produced the cell line expressing GFP::IFT52, acquired the still images of mNG::IFT81 cell line and assembled the figures. S. Bonnefoy developed the PFA/methanol fixation protocol and performed immunofluorescence data and acquisition. J. Jung characterised the mNG::IFT81 expressing cell line, performed several kymograph analyses and contributed to the setting up the High-NA microscopy in Pasteur. M. Lemos acquired transmission electron microscopy images and was responsible for measurement of the diameter of the axoneme. S. Marco contributed to statistical analysis. S. Vaughan participated to the coordination of the 3-D electron microscopy project. S. Trépout developed the scripts to perform the measurements of IFT trains (length and distance with the microtubule doublets) from FIB-SEM data and performed the corresponding statistical analysis to identify train sub-populations. J.Y. Tinevez designed the live imaging project performed at the Janelia Farm Research Campus and participated to the high-resolution video acquisition and quantification. P. Bastin conceived and coordinated the project and wrote the paper with contributions from all authors.

The authors declare no competing financial interests.

## Abbreviations

FIB-SEM: Focused Ion Beam Scanning Electron Microscopy
IFA: immunofluorescence assay
IFT: intraflagellar transport
PFR: paraflagellar rod
TEM: transmission electron microscopy

## Legends for supplementary figures

**Fig. S1.**
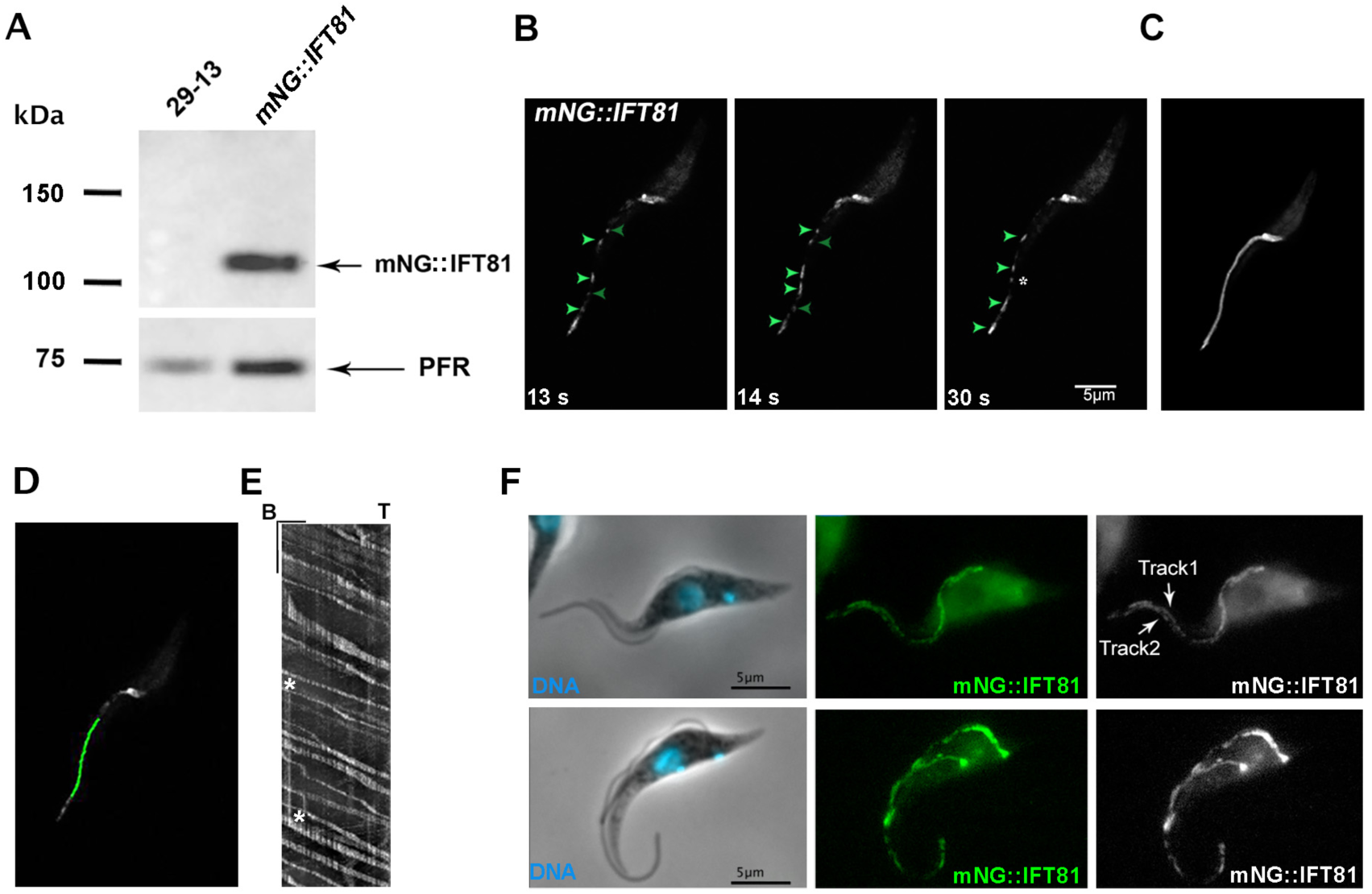
Endogenous tagging of IFT81 with mNG provides a clean marker for monitoring IFT. **(A)** Western blotting with an anti-mNG antibody reveals the expected mobility on SDS-PAGE. No signal is detected with the 29-13 (Wirtz et al., 1999) control cell line. The L13D6 monoclonal antibody recognising the PFR proteins was used as loading control. **(B)** Still images of Video S3 showing anterograde (green arrowheads) and retrograde (dark green arrowheads) IFT trafficking in a uniflagellated cell using conventional light microscopy. Focusing was made on the flagellum and the base of the flagellum is not in the same plane. This cell is growing a new flagellum that is very short and partially overlaps with the mature flagellum. The star shows an arrested IFT train. **(C)** Temporal projection shows only one track in these imaging conditions. **(D-E)** A region of interest was drawn on the indicated position of the image (D) and the kymograph was extracted showing typical robust anterograde and more discrete retrograde trains (E). The position of the arrested train marked in B is highlighted with stars. Scale bars are 2.5 µm for length (horizontal bar) and 2.5 seconds for time (vertical bar). **(F)** Cells expressing mNG::IFT81 were fixed using the paraformaldehyde-methanol protocol and direct imaging of mNG::IFT81 fluorescence was carried out revealing the existence of two parallel lines in mature and growing flagella. The base of both types of flagella is clearly visible on the bottom image.

**Fig. S2.**
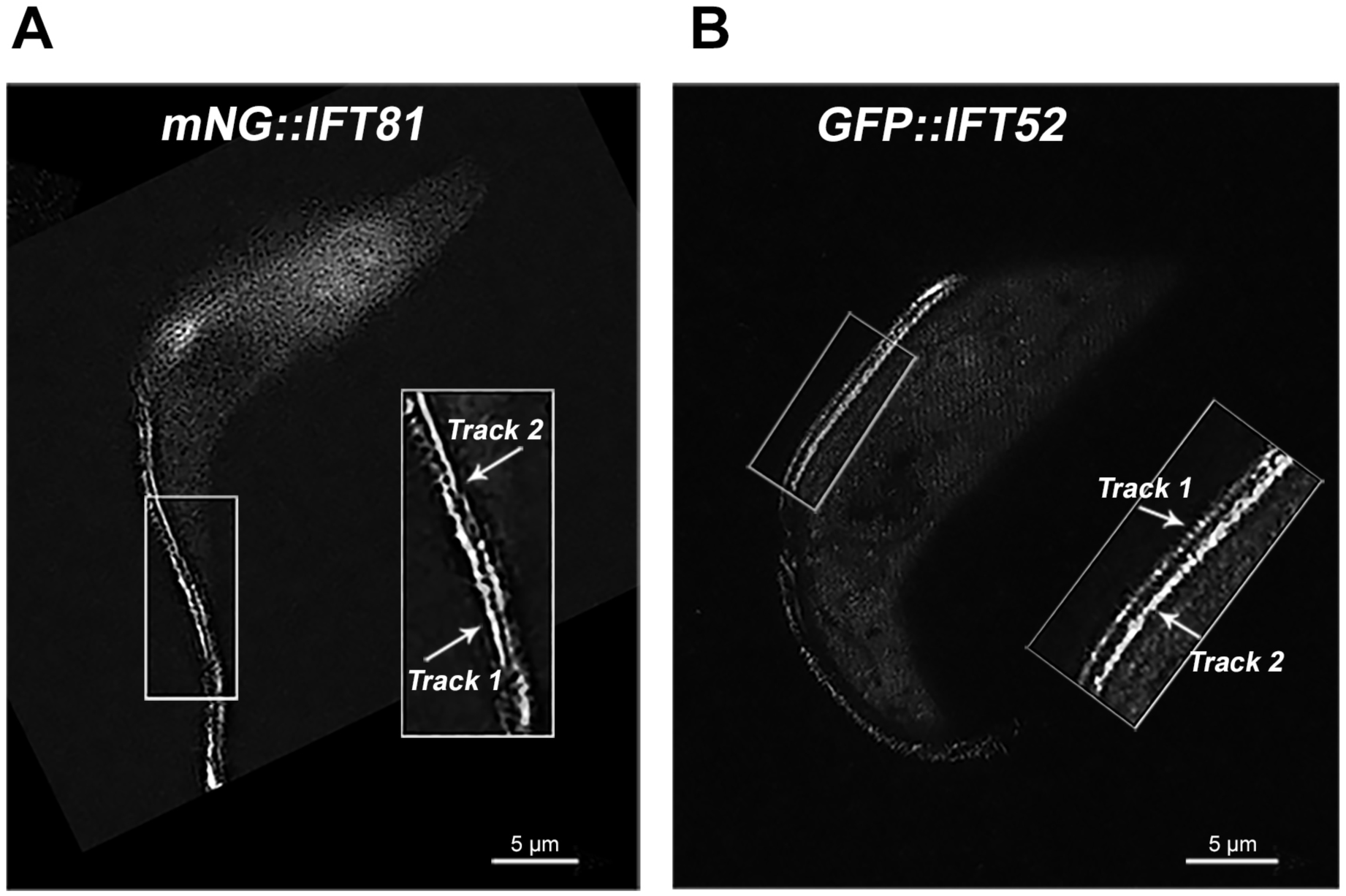
SIM imaging in live cells reveals trafficking on two separate sides of the axoneme. **(A)** SIM images showing the presence of IFT trains on two separate sides of the axoneme in cells expressing the mNG::IFT81 fusion protein. Fifteen images per plane per channel (five phases, three angles) were acquired to perform the SIM image. **(B)** Temporal projection of images coming from Video S4 in cells expressing the GFP::IFT52 fusion protein showing the presence of IFT trains on two sides of the axoneme. Areas in rectangles have been zoomed to show the two tracks. However, temporal resolution is not sufficient to determine the orientation of trafficking.

**Fig. S3.**
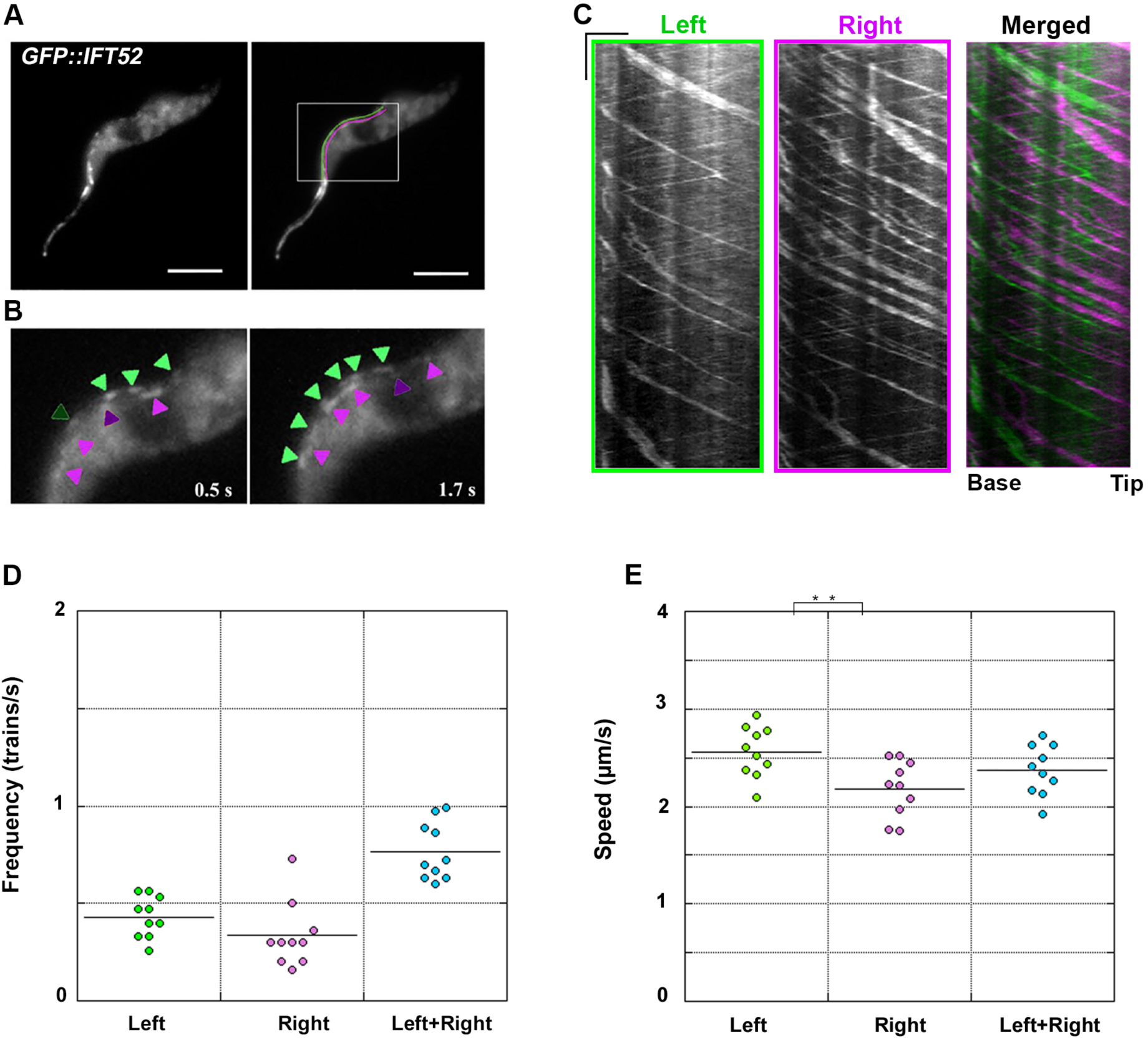
Bidirectional IFT trafficking takes place on two sides of the axoneme in live trypanosomes expressing GFP::IFT52. **(A)** Individual images from Video S6 showing the presence of two traces for IFT in the flagellum. The IFT pool at the base is out of the plane of focus. The left (green) and right (magenta) sides were defined after orientating the cell with the posterior end on top of the image and the flagellum on the left-hand side. **(B)** Still images from Video S6 of live trypanosomes expressing GFP::IFT52 imaged at high-resolution. Green and magenta arrowheads indicate trains on the left and right tracks with lighter arrowheads pointing at anterograde trains and darker arrowheads showing retrograde trains. **(C)** Kymograph analysis of the same cell showing the left (green), the right (magenta) and the merged images. Scale bars are 2 µm for length (horizontal bar) and 2 seconds for time (vertical bar). **(D)** Dot plot of the frequency of anterograde IFT trains visible on the left (green) and the right (magenta) side and the sum of both (cyan). **(E)** Dot plot but for the speed of anterograde trains.

**Fig. S4.**
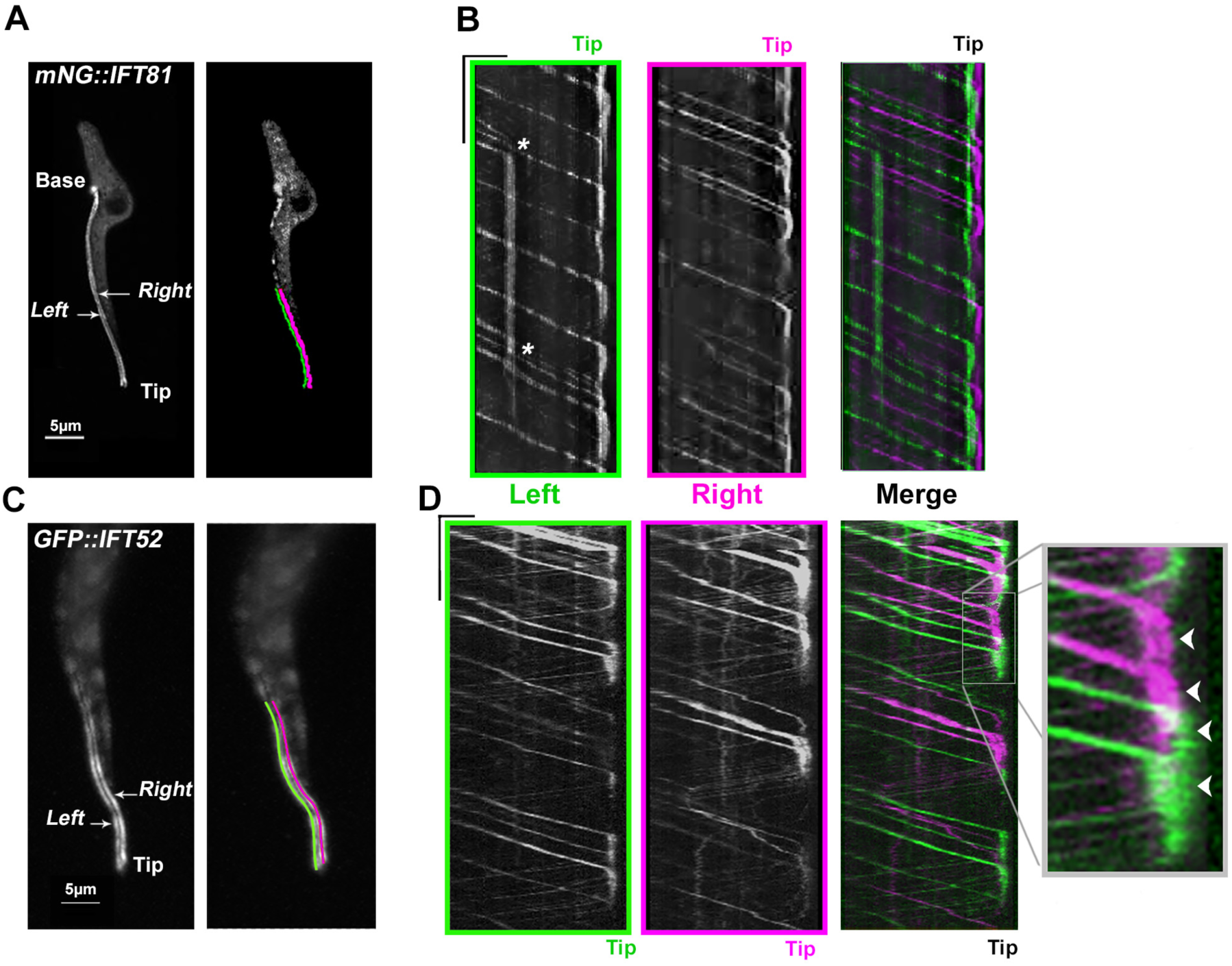
Anterograde trains are converted to retrograde trains after a transition period at the distal tip. **(A)** Temporal projection of the flagellum in a cell expressing mNG::IFT81. The region of interest at the tip of the flagellum is indicated, with left (green) and right (magenta) **sides of the axoneme**. See corresponding Video S9. **(B)** The kymographs are shown for each of them is the corresponding colour. At the distal end, IFT proteins present in anterograde trains transit for a few seconds before returning in smaller and less bright retrograde trains. Note the presence of an anterograde train that arrested for a few seconds before the material it contained was picked up by other anterograde trains (stars). Scale bars are 2.5 µm for length (horizontal bar) and 5 seconds for time (vertical bar). **(C)** The distal tip of a cell expressing GFP::IFT52 was photobleached and the transition from anterograde to retrograde trains was monitored in the indicated region of interest. **(D)** Kymographs are highlighted in green and magenta as above. Scale bars are 2.5 µm for length (horizontal bar) and 5 seconds for time (vertical bar). The enlarged portion shows the distal end of the flagellum where fluorescent proteins present in anterograde trains are seen transiting for a few seconds (indicated by white arrowheads) before leaving in association to multiple but discrete retrograde trains. No evidence for transfer between the left and rights tracks could be observed.

**Video S1**. Three-dimensional view of wild-type trypanosomes analysed by FIB-SEM with the stack of original data where several cells are visible with all typical organelles including the flagellum. **The position of three IFT trains present in the flagellum of the cell located in the centre of the field is indicated with arrowheads. The video pauses each time a new IFT train has been annotated. The same cell has been used for the segmentation analysis presented at Figure 3. The same reference number has been used for these IFT trains (IFT2, IFT3 and IFT4). To reduce the size of the video, only a portion of the stack is shown. It does not contain the full flagellum explaining why the train labelled IFT1 is not visible.**

**Video S2**. Individual flagella of the stack from Video S1 are shown in different colours and the IFT trains are shown in red. The volume is then rotated in all dimensions to visualise the positioning of IFT trains.

**Video S3**. Live imaging of a cell expressing mNG::IFT81 showing robust bidirectional IFT.

**Video S4**. Live imaging by SIM of a cell expressing GFP::IFT52 showing the existence of two tracks for IFT trafficking. Time-series were acquired for a total of 14 seconds. Although the spatial resolution allows the distinction of two tracks, the time resolution is not sufficient to discriminate individual IFT trains on them.

**Video S5**. Live imaging by high-resolution microscopy using a 1.57 NA objective of a cell expressing mNG::IFT81. Bidirectional IFT trafficking is visible on two distinct lines. The spatial resolution allows the distinction of two sides of the axoneme and the time resolution permits the discrimination of individual IFT trains on each of them.

**Video S6**. Live imaging by high-resolution microscopy using a 1.49 NA objective of a cell expressing GFP::IFT52. Bidirectional IFT trafficking is visible on two distinct lines. The spatial resolution allows the distinction of two sides of the axoneme and the time resolution permits the discrimination of individual IFT trains on each of them.

**Video S7**. **Live imaging by high-resolution microscopy using a 1.57 NA objective of a cell expressing GFP::DHC2.2. Bidirectional IFT trafficking is visible on two distinct lines. The spatial resolution allows the distinction of two sides of the axoneme and the time resolution permits the discrimination of individual IFT trains on each of them.**

**Video S8**. **Live imaging by conventional microscopy using a 1.40 NA objective of a cell expressing mNG::IFT81 (green) and TdT::DHC2.1 (magenta). The two proteins co-localise on moving and standing trains (see kymograph analysis at Figure 6C).**

**Video S9**. Live imaging by high-resolution microscopy using a 1.57 NA objective of a cell expressing mNG::IFT81. Focusing on the distal tip reveals the transit and turnaround of IFT material during the conversion of anterograde to retrograde trains.

**Video S10**. **Live imaging by high-resolution microscopy using a 1.49 NA objective of a cell expressing GFP::IFT52. Focusing on the distal tip reveals the transit and turnaround of IFT material during the conversion of anterograde to retrograde trains. For technical reasons, the bleaching event could not be recorded and only the recovery phase can be presented. See Figure 7 for detailed legend.**

